# Comprehensive chromatome profiling identifies metabolic enzymes on chromatin in healthy and cancer cells

**DOI:** 10.1101/2023.12.06.570368

**Authors:** S Kourtis, M Guirola, N Pardo-Lorente, R Ghose, M Garcia-Cao, A Gañez-Zapater, S Haynes, F Fontaine, A Muller, S Sdelci

## Abstract

Metabolic and epigenetic rewiring are widely considered hallmarks of cancer, with emerging evidence of crosstalk between them. Anecdotal evidence of metabolic enzymes moonlighting in the chromatin environment has suggested how this crosstalk might be facilitated, but the extent of nuclear relocalization of metabolic enzymes remains elusive. Here, we provide a comprehensive chromatin proteomics resource across cancer lineages as well as healthy samples and demonstrate that metabolic enzyme moonlighting on chromatin is widespread across tissues and pathways. We show that the abundance of metabolic enzymes on chromatin is tissue-specific, with oxidative phosphorylation proteins depleted in lung cancer samples, perhaps suggesting an interplay between cell identity and nuclear metabolism. Finally, we explore metabolic functions in the chromatin environment and show that one-carbon folate enzymes are associated with DNA damage and repair processes, providing an approach to explore non-canonical functions of metabolic enzymes.

## Introduction

While broad changes in the epigenetic and metabolic landscapes are both recognized hallmarks of cancer (De Berardinis & Chandel, 2016; Flavahan et al., 2017; Pavlova & Thompson, 2016), their interplay has remained elusive. During cancer transformation, a complex metabolic rewiring is required to sustain tumor development and thereby support the metastatic process. Canonically, the interplay between metabolism and epigenetics is mediated by the availability of metabolic substrates, such as acetyl-CoA, S-adenosylmethionine (SAM), or adenosine triphosphate (ATP), which facilitate epigenetic modifications (Gapa et al, 20-22., 2022; Kinnaird et al., 2016; Lu & Thompson, 2012; Wong et al., 2017)), which could lead to transcriptional changes of specific metabolic enzymes (Dang, 2013; Morandi et al., 2017). While classic examples such as the c-myc oncogene transcriptionally promoting folate metabolic enzymes (Morandi et al., 2017) and mutations in isocitrate dehydrogenase 1 and 2 (IDH1 and IDH2) associated with aberrant genome-wide DNA and histone hypermethylation (Yan et al., 2009) demonstrate the indirect relationship between epigenetics and metabolism, evidence for a more direct interaction is accumulating (Yang et al., 2022).

Enzymes with nuclear moonlighting functions include the alternative isoform of the enolase enzyme (ENO-1), which localizes to chromatin and competes with c-myc, thereby regulating its activity (Didiasova et al,., 2019), GAPDH induces apoptotic gene expression (Sirover, 2021), nuclear FBP1 interacts with the polycomb repressive complex (PRC2) to regulate levels of H3K27me3 (Bian et al., 2022), and finally nuclear PKM2 phosphorylates histone H3(T11) to regulate histone acetylation (Gupta & Uversky, 2023; Jeffery, 2019). In addition, the nuclear enzymatic activity of certain metabolic enzymes has also been demonstrated. Nuclear-generated acetyl-CoA has been shown to directly influence histone acetylation (Bulusu et al., 2017), and nuclear-generated ATP in breast cancer cells is essential for estrogen-induced chromatin remodeling (Wright et al., 2016). Finally, we have previously shown that MTHFD1, a predominantly cytoplasmic folate metabolism enzyme, is present on chromatin and is a direct interactor and activity modulator of the epigenetic reader BRD4 (Sdelci et al., 2019). The in loco product of folate metabolites on chromatin, facilitated by this interaction, was necessary to maintain the transcriptional profile of the cell, as they could be immediately used for nucleotide synthesis. Thus, the presence and activity of metabolic enzymes on chromatin could be widespread and an integral part of epigenetic regulation.

With the goal of comprehensively identifying novel moonlighting or non-canonical nuclear functions of metabolic enzymes, we have optimized a subcellular fractionation protocol that enriches chromatin-bound proteins across a panel of tissue types for both cancerous and untransformed cells. In addition to canonical chromatin proteins, we identify metabolic enzymes that consistently localize to chromatin, many with tissue- and cancer-specific localization. We uncover metabolic pathways that are present on chromatin, such as oxidative phosphorylation and one-carbon by folate, and explore their functional relationships on chromatin.

## Results

To achieve characterization of chromatin-bound proteins across tissue types, we optimized the previously published chromatin enrichment protocol (Sdelci et al., 2019) for maximum chromatin enrichment across 10 tissue types (Fig. 1A, Table 1).We analyzed a total of 108 samples, including 44 cancer cell lines and 10 untransformed cells, in duplicate, which represents a very extensive sample collection. This allowed retrieval of chromatin-enriched samples, for both established cancer cell lines but also their untransformed counterparts (Fig. 1B). We visually followed the fractionation protocol, to ensure that non-nuclear proteins (FDX1) were sequentially depleted from the samples during the washes, before the nuclear lysis step (Fig. 1C, Appendix 1). Using focal adhesion (Vinculin), mitochondrial (FDX1) and histone markers (H3), we validate that the chromatin-bound proteomic samples were indeed depleted of non-chromatin markers and enriched for histones compared to the cytoplasmic fraction (Fig. 1D, Appendix 2). Samples were acquired using Data-Independent Acquisition Mass spectrometry (DIA-MS) and showed a strong enrichment for nuclear and chromosome-related terms and depleted mitochondrial and cytoplasmic proteins (Fig. S1A), further validating the protocol and MS acquisition. We achieve approximately 5100 proteins per sample, most quantified by multiple peptide-to-spectrum matches (PSMs), with high reproducibility in the number of retrieved proteins between biological replicates (Fig. 1E). While most proteins were identified and quantified across the majority of samples (Fig. S1B), some proteins were only sporadically detected (Table 2). We speculate that proteins which are sparsely present in our dataset are enriched for sample-specific non-chromatin proteins. To demonstrate this, we predicted sequence-based nuclear localisation signals (NLS) genome-wide and use them as a proxy for nuclear proteins (Fig. S1C) due to their agreement with known nuclear proteins (Cokol et al., 2000; Mulvey et al., 2017). Indeed, proteins with very few missing values were enriched for known nuclear proteins and proteins sparsely detected in the data were depleted for nuclear proteins. As such we defined a core-chromatome, comprising of 3467 proteins which were consistently found on chromatin across the majority of tissue types. Among other proteins, we retrieve well-known chromatin complexes (Fig. 1F) such as SWI/SNF and HDAC proteins as ‘core-chromatome’ proteins (Drew et al., 2021; Huttlin et al., 2017).

**Figure 1:**
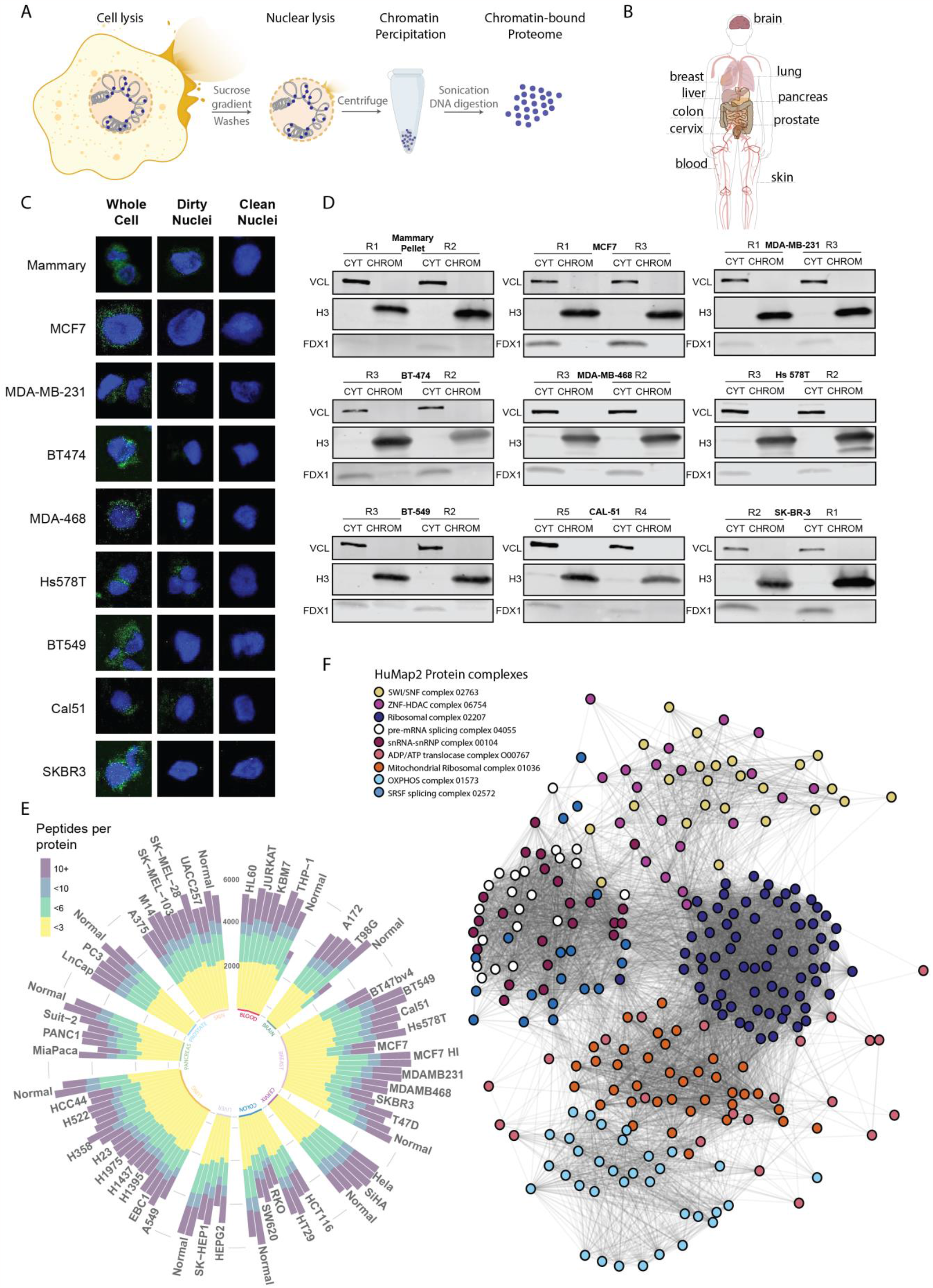
Chromatin enrichment protocol identifies proteins on chromatin across cancer lineages a. Protocol of chromatin enrichment for proteomic profiling b. Tissue types included in chromatin proteome characterization c. Immunofluorescence images of protocol progression to achieve enriched chromatin of breast samples with DAPI (blue) and FDX1 (green) shown. d. Western blots of cytoplasmic and chromatin enriched samples with non-nuclear (Vinculin), chromatin (Histone 3) and mitochondrial markers (FDX1) across breast samples. e. Total number of detected proteins, with their respective PSMs per samples, acquired by DIA-MS. f. Known hu.MAP 2.0 complexes detected in chromatin samples, with edges representing known protein-protein interactions from BioPlex 2.0.

To identify functional pathways on chromatin, we searched the core-chromatome for enriched ‘Biological Processes’ gene sets. We retrieved many well-characterized processes on chromatin such as ‘DNA repair’ and ‘DNA replication’, as well as surprisingly ‘ATP metabolic process’ (Fig. S2A). This term was mainly driven by proteins that constitute the electron transport chain (ETC), such as members of the NDUF family. To check which other metabolic pathways were present on chromatin, we repeated the analysis specifically against KEGG pathways (Kanehisa et al., 2023; Kanehisa & Goto, 2000). Among other pathways, oxidative phosphorylation was found to be the most complete pathway on chromatin, with more than 60% of subunits we identify in our dataset, being consistently present on chromatin (Fig. 2A, Fig. S2B). Oxidative phosphorylation was followed by Lysine degradation, which includes lysine methyltransferase family members that are well established to be on chromatin, such as the KMTs, NSDs and SETDs. To explore tissue-specific pathways on chromatin, we annotated core-chromatome protein using KEGG pathways and clustered proteins based on their detection in each sample. We observed a strong tissue-based clustering of samples, with blood, lung, skin, brain and cervix showing high consistency across cell lines, while breast and pancreas demonstrated high heterogeneity in the chromatin bound proteome across cell lines (Fig. S2C). Interestingly, proteins belonging to the ‘Thermogenesis’ term, were consistently detected across samples, with the exception of lung, colon and pancreas samples, suggesting that these proteins were absent from the chromatin environment in these tissues. ‘Thermogenesis’ consists primarily of proteins related to the oxidative phosphorylation pathway, which prompted us to look specifically for metabolic proteins in our dataset, since most proteins with enzymatic function canonically exist outside the nucleus. Interestingly, while lung samples were depleted in oxidative phosphorylation protein, some breast samples appear to be depleted in pyrimidine and pyruvate metabolism proteins (Fig. 2B), suggesting tissue-specificity.

**Figure 2:**
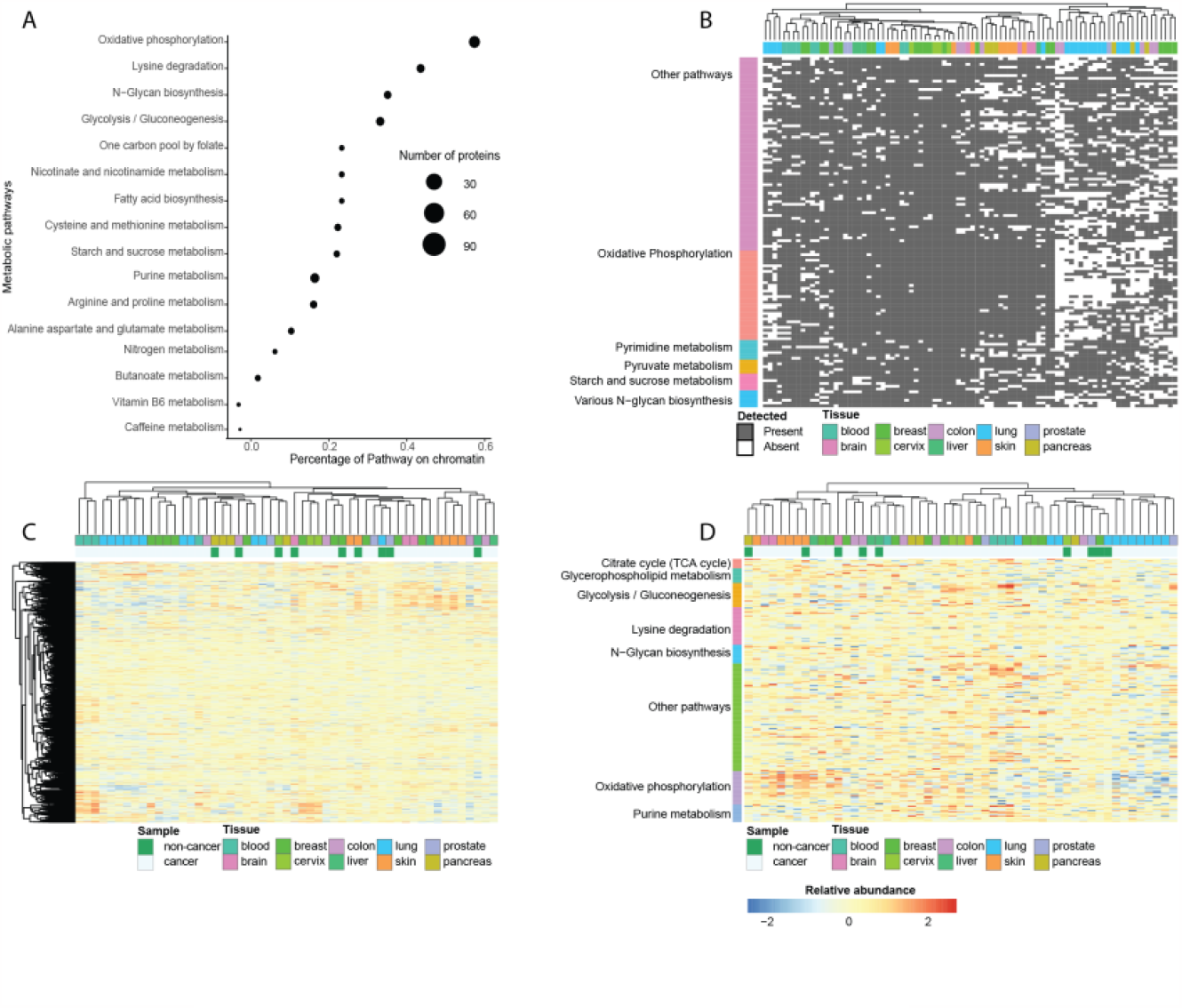
Metabolic pathways on chromatin a. Percentage of pathway detected on chromatin for most complete and incomplete KEGG pathways. b. Clustering of chromatin proteomic samples based on detection of metabolic KEGG enzymes. c. Clustering of chromatin proteomics samples based on relative intensity of all proteins detected in the samples. d. Clustering of chromatin proteomics samples based on relative intensity of KEGG enzymes detected in the samples.

We observed a variation in the degree of chromatin enrichment, as quantified by the intensity of total histone proteins across samples (Fig. SF2D), likely due to subtle differences in cell-line specific properties, causing variation in the cell lysis and chromatin enrichment protocol. To make the samples comparable for downstream analysis, we regressed all protein abundances against the relative enrichment of each sample (Fig. S2E), where we expected nuclear proteins to have a positive coefficient against chromatin enrichment and non-nuclear proteins a negative coefficient. This is demonstrated by a pair of primarily non-nuclear (LAMP2) and nuclear (H3-3) proteins(Thul et al., 2017), which behaved differently relative to the enrichment term, as a consequence of their relative proportion on chromatin (Fig. 2F). This regression removed the compartment-specific separation within our dataset (Fig. 2G-H), with the largest differences on chromatome abundances being present between blood and skin cell lines (Fig. 2I). Further confirming our model, regression coefficients of each protein, representing relative protein proportion on chromatin, strongly agreed with hyperLOPIT subcellular annotations and the presence of NLS motifs for nuclear proteins (Fig. S2J-K). Quantitatively, we observed blood, skin, lung, and pancreas tissues to display more homogeneity regarding their chromatin-bound proteins, with the non-transformed pancreas sample showing the closest similarity to its respective tissue (Fig. 2C). Focusing on metabolic pathways, we observed that enzymes on chromatin vary in a tissue-specific manner, leading to the clustering of cancer types (Fig. 2D). While skin and brain showed high levels of chromatin-bound oxidative phosphorylation proteins compared to lung samples, blood samples exhibited higher levels of purine metabolism proteins (Fig. 2D). Although many of these enzymes show the ability to re-localize to chromatin, their abundance appeared to be tissue dependent. Interestingly, most of these enzymes are smaller than the ∼80kDa nuclear pore, with only a small fraction of their total pool on chromatin (Fig. S2L). Proteins with enzymatic functions containing NLS signals were the exception, with their canonical localization being nuclear, such as KMTs with a high fraction on chromatin, or another membrane-bound compartment, such as ALG3 and STT3B with only a fraction on chromatin. Finally, enzymes larger than 80 kDa and lacking an NLS signal do not seem to be excluded from chromatin, ruling out the possibility that their nuclear localization arises by passive diffusion (Fig. 2L).

Among proteins which we identified consistently on chromatin, 335 proteins were differentially present in at least one cancer type, compared to its non-cancerous counterpart (Fig. 3A). Surprisingly, metabolic enzymes, additionally to being present on chromatin in a tissue-specific manner, also seemed to be differentially abundant in cancer. For instance, Acyl-CoA-binding domain-containing protein 5 (ACBD5) was significantly more abundant on chromatin in liver cancer, while depleted in cervix and prostate cancer (Fig. 3B, upper). Similarly, ATP synthase membrane subunit j (ATP5MJ), was more abundant in cervix cancer and depleted from brain and skin cancers (Fig. 3B, lower). While both proteins showed nuclear localisation, further supporting our data of oxidative phosphorylation subunits being in the nucleus, there was no previous characterization of their nuclear function (Fig. 3C).

**Figure 3:**
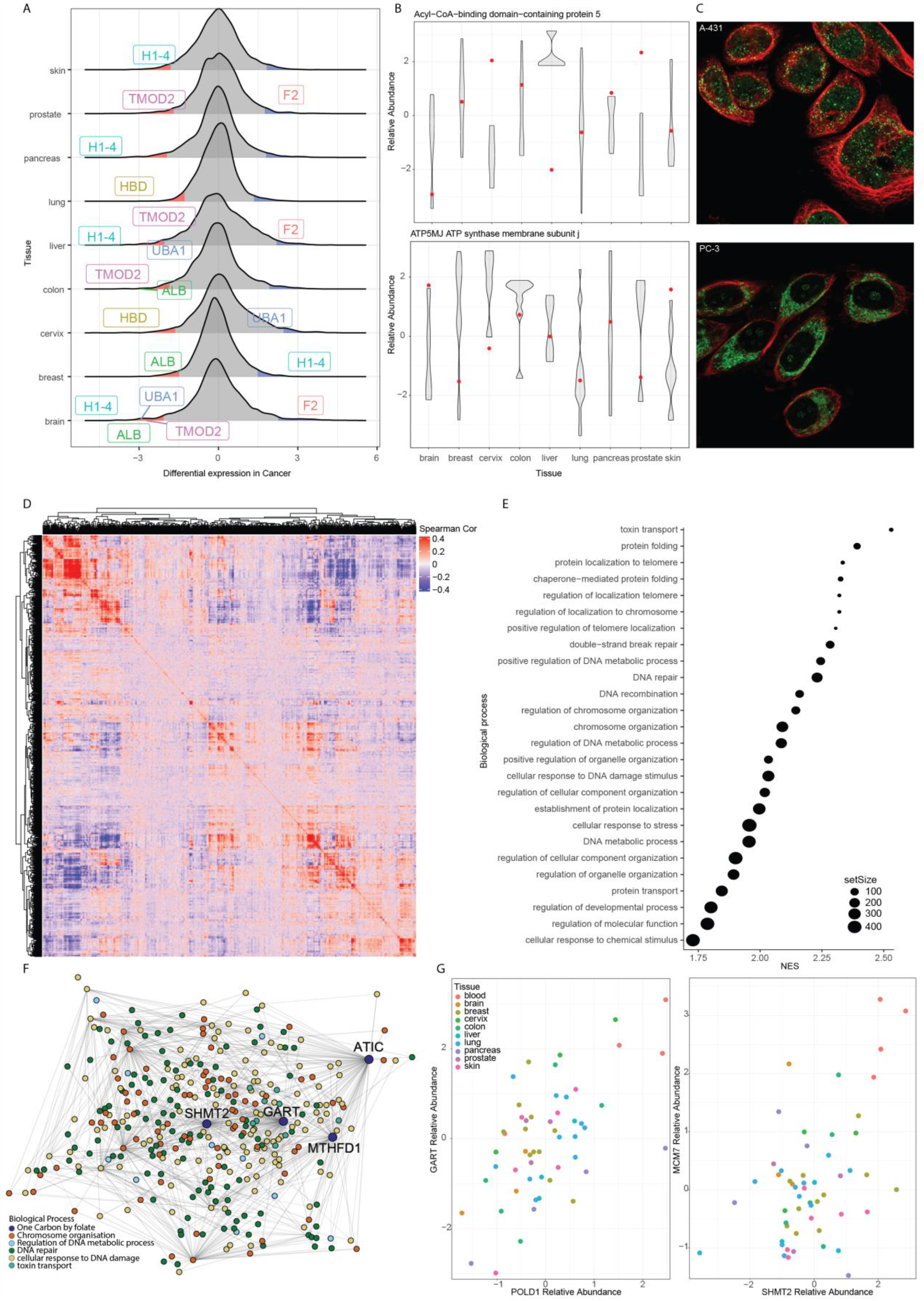
Differentially abundant proteins on chromatin a. Differentially present proteins on chromatin per cancer lineage, compared to healthy counterpart samples. b. ACBD5 and ATP5MJ protein abundance in chromatin samples, based on lineage, with abundance in healthy counterpart represented in red. c. Immunohistochemistry images in PC-3 and A-431 cells of ATP5MJ and ACBD5 from HPA. d. Protein correlation structure across chromatin samples. e. Biological process enrichment of proteins correlated to one-carbon by folate enzymes found in chromatin. f. Protein correlation of pathways significantly correlated with one-carbon by folate pathway enzymes. g. Relative protein abundance per sample for GART-POLD1 and MCM7-SHMT2 protein pairs.

To elucidate protein function on chromatin, we explored protein covariation in our data, following previous studies showing that proteins that covary are functionally related (Kustatscher et al., 2019). Proteins on chromatin displayed functional patterns, with highly correlated proteins in our dataset annotated as interactors in covariation studies (Fig. 3D, Fig. F3A). Examples of proteins highly covarying on chromatin tended to belong to chromatin complexes, which is expected since complex members are known to be highly regulated at the protein level (Fig. S3B). Using the covariation structure of our data, we discovered that enzymes belonging to the ‘one carbon by folate’ pathway, when on chromatin, are functionally related to telomere localization, DNA metabolism, and DNA damage and repair (Fig. 3E-F). To disentangle whether these proteins are related in a tissue-specific or pan-cancer manner, we explored these correlations. Whereas Trifunctional purine biosynthetic protein adenosine-3 (GART) and DNA polymerase delta catalytic subunit (POLD1) correlated across tissues, the relationship between Serine hydroxymethyltransferase (SHMT2) and DNA replication licensing factor MCM7 (MCM7) was mainly driven by their high abundance in blood chromatin samples.

An alternative approach to understanding protein function is to study gene essentiality and drug targeting, where proteins that are similarly affected by drugs or covary in essentiality are functionally related (Dempster et al., 2021). By correlating our chromatin abundance data with publicly available gene essentiality and drug sensitivity data, we revealed the expected positive correlation between BANP abundance, which promotes TP53 activation, and sensitivity to the TP53-MDM2 inhibitor nutlin-3 (Fig. S3C, top). Interestingly, we also observe that CDKN1A interacting zinc finger protein 1 (CIZ1) chromatin abundance correlated with SRC kinase signaling inhibitor 1 (SRCIN1) gene dependency (Fig. 3C, bottom). While SRCIN1 is canonically located at the cell periphery, its nuclear localization may indicate a moonlighting nuclear function responsible for the functional relationship with CIZ1 (Fig. 3F, bottom, S3D).

## Discussion

While moonlighting of specific metabolic enzymes has long been recognized, recent evidence indicates that nuclear moonlighting is more widespread than previously thought. Here, we provide a systematic approach to identify proteins on chromatin in healthy and cancerous tissues, revealing the universality of nuclear moonlighting of enzymes across lineages. Surprisingly, we discover tissue- and cancer-specific enzymes on chromatin, suggesting an interplay between cell identity and nuclear metabolism. Among others, oxidative phosphorylation enzymes appear to be collectively present on chromatin, supporting our previous observations (Moretton et al., 2023), while surprisingly absent from specific tissues such as lung and colon. While exploring the moonlighting functional role of proteins, our results identified folate enzymes associated with DNA damage and repair, confirming at the pathway-level our previous characterization of chromatin-bound MTHFD1 (Sdelci et al., 2019). Collectively, our results contribute to the growing literature on the presence and functionality of nuclear metabolism, while demonstrating that enzymes on chromatin exhibit lineage-specific distributions that may suggest an interplay between cell identity and nuclear metabolism.

## Methods

### Cell culture

All cell lines were cultured at 37°C in 5% CO2. The culture conditions are specified in Supplementary Table 1. Cell cultures were tested every month for mycoplasma contamination.

### Chromatome fractionation

To obtain the chromatome samples for immunofluorescence validation, 2.5^10^6^ cellular pellets were first resuspended in 75 uL of PBS. Immediately, a 15 uL aliquot (whole-cells) was taken and fixed in fixative solution (methanol 100%) for later immunofluorescence. 75 uL of CHAPS (3-cholamidopropyl dimethylammonium 1-propane sulfonate) Buffer was added to the sample to break the cytosolic membrane. The concentration and time of lysis for each cell line is specified in Supplementary Table X. Cell lysates were checked at the microscope to verify the correct cell lysis. Then, lysed cells were centrifuged for 5 min at 720g at 4ºC and the supernatant was discarded. The nuclear pellet was resuspended in Cytoplasmic Lysis Buffer (IGEPAL 0.1%, NaCl 150 mM, Tris-HCl 10 mM pH 7 in H2O), where a 100 uL aliquot (dirty nucleus) was taken and fixed in fixative solution for later immunofluorescence. The rest of the sample was placed on the top of a Sucrose Gradient Buffer (NaCl 150 mM, sucrose 25%, Tris-HCl 10 mM pH 7 in H2O) and centrifuged for 15 min at 10,000g at 4ºC. Purified nuclei were then washed 3 times by resuspending in Nuclei Washing Buffer (EDTA 1 mM, IGEPAL 0.1% in PBS) and centrifuged for 5 min at 1200g at 4ºC. Finally, the resulting sample (clean nuclei) was resuspended in fixative solution for later immunofluorescence.

To obtain the chromatome samples for mass spectrometry, 5^10^6^ cellular pellets were first resuspended in 150 uL of CHAPS (3-cholamidopropyl dimethylammonium 1-propane sulfonate) Buffer to break the cytosolic membrane. The concentration and time of lysis for each cell line is specified in Supplementary Table X. Then, lysed cells were centrifuged for 5 min at 720g at 4ºC and the supernatant was harvested as the cytosolic fraction. The nuclear pellet was resuspended in Cytoplasmic Lysis Buffer (IGEPAL 0.1%, NaCl 150 mM, Tris-HCl 10 mM pH 7 in H2O) and placed on the top of a Sucrose Gradient Buffer (NaCl 150 mM, sucrose 25%, Tris-HCl 10 mM pH 7 in H2O) and centrifuged for 15 min at 10,000g at 4ºC. Purified nuclei were then washed 3 times by resuspending in Nuclei Washing Buffer (EDTA 1 mM, IGEPAL 0.1% in PBS) and centrifuged for 5 min at 1200g at 4ºC. Then, the washed nuclear pellet was resuspended in Nuclei Resuspension Buffer (EDTA 1 mM, NaCl 75 mM, 50% sucrose, Tris-HCl 20 mM pH 8 in H2O) and the nuclear membrane was lysed by adding Nuclei Lysis Buffer (EDTA 0.2 mM, HEPES 20 mM pH 7.5, IGEPAL 0.1%, NaCl 300 mM in H2O), vortexing and incubating for 5 min. After centrifugation for 2 min at 16000g at 4ºC, the resulting chromatin was resuspended in Benzonase Digestion Buffer (15 mM HEPES pH 7.5, 0.1% IGEPAL, TPCK 5 mg/mL) and sonicated on a Bioruptor Pico (Diagenode) for 15 cycles 30 sec ON/30 sec OFF in 1.5 mL Diagenode tubes (Diagenode; #C30010016). Finally, sonicated chromatin was digested with benzonase enzyme (VWR; #706643; 2.5U) for 30 min at room temperature, and the resulting sample was harvested as chromatome fraction. All the steps were performed on ice and all buffers were supplemented with proteinase inhibitors (Roche; #4693132001). Cytosolic and chromatome extracts were quantified with Pierce BCA Protein Assay Kit (Thermo Scientific; #PIER23225).

### Immunofluorescent imaging of chromatome samples to test validity

Samples collected at unlysed cell, pre-nuclear wash and post-nuclear wash phases of the fractionation protocol were used to perform IF to validate the integrity of the fractions. Briefly, samples were fixed in methanol overnight and IF performed in black 96-well plates for microscopy. Fixed samples were spun down at 1000 RCF and resuspended in PBS. Each sample was then placed in one well of the 96-well plate and the plate was centrifuged at 4oC, 1500 rcf for 15 min to attach the cells to the bottom. The PBS was then gently aspirated using a pipette and samples incubated in TBP (0.1% Triton-X-100 in 2% BSA/PBS) for 30 min at room temperature (RT). TBP was then aspirated gently using a pipette and samples incubated with the primary antibodies. Staining for mitochondria was done using 30 ul of 1:200 dilution (in 0.5% BSA/PBS) FDX-1 antibody (Thermo Fisher Scientific, PA559653) for 1 hour at RT. Samples were washed twice with PBS and further incubated with 30uL of 1:200 Alexa Fluor 488 secondary antibody in 0.5% BSA/PBS. Following two washes with PBS, nuclear staining was done using 1:1000 dilution (in PBS) of DAPI (DAPI ref) for 5 min at RT. Following a PBS wash, samples were then imaged.

Imaging was done using confocal microscopy on the LEICA TCS SP5 inverted microscope in collaboration with the Advanced Light Microscopy Unit (ALMU) at CRG, Barcelona. This microscope is equipped with Leica inverted confocal system (405, 458, 476, 488, 496, 514, 561 and 633 nm lasers). The 405 laser and the 488 laser were used for excitation of DAPI (Ex: 405 nm /Em: 461 nm) and FDX1 (Ex: 488 nm /Em: 515 nm). Image acquisition was performed under oil immersion at 63x magnification. Visualisation and figure assimilation was done with the help of ImageJ (Schindelin et al., 2012; Schneider et al., 2012).

### Western Blot

Cytosolic and chromatin fractions were quantified by Micro BCA™ Protein Assay Kit (Thermo Fisher Scientific #23235) according to manufacturer instructions. Proteins were resolved on 4–20% gradient precast Mini-PROTEAN TGX polyacrylamide gels (BIO-RAD #4561094) and transferred to a 0.45 mm nitrocellulose blotting membrane (Amersham™ #10600002) for immunoblotting. Primary antibodies used were Vinculin (E1E9V) XP^®^ rabbit monoclonal antibody (Cell Signaling Technology #13901; 1/1000), FDX1 rabbit polyclonal antibody (Thermo Fisher Scientific # PA5-59653, 1/1000) and Histone H3 (1B1B2) mouse monoclonal antibody (Cell Signaling Technology #14269, 1/10000). Fluorescence-conjugated secondary antibodies Alexa Fluor™ Plus 800 goat anti-rabbit IgG (Thermo Fisher Scientific; #A32735; 1:10000); Alexa Fluor™ Plus 800 goat anti-mouse IgG (Thermo Fisher Scientific; #A32730; 1:10000); Alexa Fluor™ 680 goat anti-mouse IgG (Thermo Fisher Scientific; #A21058; 1:10000) and Alexa Fluor™ Plus 680 goat anti-rabbit IgG (Thermo Fisher Scientific; #A32734; 1:10000) were used for signal detection with Odyssey CLx Imaging System (LI-COR Biosciences).

### Data Analysis

Data processing: Chromatin data were analysed using DIA-NN software (Demichev et al., 2020) which were subsequently normalized using the normalize_vsn and median_normalisation functions from the DEP (Zhang et al., 2018) and proDA (Ahlmann-Eltze, 2022) packages, respectively (R Core Team, 2022). The rest of the pipeline was followed according to the DEP package, with the inclusion of impute.mi function for protein-imputation from the imp4p package (Gianetto, 2021). Relative enrichment of each sample was estimated as the relative abundances of known histone proteins in each sample (Mulvey et al., 2017) and all biological enrichment performed using the clusterProfiler tool (Wu et al., 2021), and regression correction was performed using the rlm function (Venables & Ripley, 2002). Known subcellular localizations for proteins were obtained from the pRoloc R package, hyperLOPITU2OS2018 (Gatto et al., 2018). Analysis was facilitated by the tidyverse (Wickham et al., 2019) and data.table (Dowle & Srinivasan, 2022) collection of packages. Essentiality and drug sensitivity data were conducted by comparing chromatin abundances to gene essentialities and drug sensitivities (Dempster et al., 2019).

### Liquid chromatography coupled to tandem mass spectrometry (LC-MS/MS)

The protein concentrations from chromatin enriched samples were determined using the BCA protein assay kit (Applichem CmBH, Darmstadt, Germany), and 10 μg per sample was processed using an adapted Single-Pot solid-phase-enhanced sample preparation (SP3) methodology (Hughes et al., 2014). Briefly, equal volumes (125 μl containing 6250 μg) of two different kind of paramagnetic carboxylate modified particles (SpeedBeads 45152105050250 and 65152105050250; GE Healthcare) were mixed, washed three times with 250 μl water and reconstituted to a final concentration of 50 μg/μl with LC-MS grade water (LiChrosolv; MERCK KgaA). Samples were filled up to 100 μL with stock solutions to reach a final concentration of 2% SDS, 100mM HEPES, pH 8.0, and proteins were reduced by incubation with a final concentration of 10 mM DTT for 1 hour at 56°C. After cooling down to room temperature, reduced cysteines were alkylated with iodoacetamide at a final concentration of 55 mM for 30 min in the dark. For tryptic digestion, 400 μg of mixed beads were added to reduced and alkylated samples, vortexed gently and incubated for 5 minutes at room temperature. The formed particles-protein complexes were precipitated by addition of acetonitrile to a final concentration of 70% [V/V], mixed briefly via pipetting before incubating for 18 minutes at room temperature. Particles were then immobilized using a magnetic rack (DynaMag-2 Magnet; Thermo Fisher Scientific) and supernatant was discarded. SDS was removed by washing two times with 200 μl 70% ethanol and one time with 180 μl 100% acetonitrile. After removal of organic solvent, particles were resuspended in 100 μl of 50 mM NH4HCO3 and samples digested by incubating with 1 μg of Trypsin overnight at 37°C. Samples were acidified to a final concentration of 1% Trifluoroacetic acid (Uvasol; MERCK KgaA) prior to immobilizing the beads on the magnetic rack. Peptides were desalted using C18 solid phase extraction spin columns (Pierce Biotechnology, Rockford, IL). Finally, eluates were dried in a vacuum concentrator and reconstituted in 10 μl of 0.1% TFA.

Mass spectrometry was performed on an Orbitrap Fusion Lumos mass spectrometer (ThermoFisher Scientific, San Jose, CA) coupled to an Dionex Ultimate 3000RSLC nano system (ThermoFisher Scientific, San Jose, CA) via nanoflex source interface. Tryptic peptides were loaded onto a trap column (Pepmap 100 5μm, 5 × 0.3 mm, ThermoFisher Scientific, San Jose, CA) at a flow rate of 10 μL/min using 0.1% TFA as loading buffer. After loading, the trap column was switched in-line with a 50 cm, 75 μm inner diameter analytical column (packed in-house with ReproSil-Pur 120 C18-AQ, 3 μm, Dr. Maisch, Ammerbuch-Entringen, Germany). Mobile-phase A consisted of 0.4% formic acid in water and mobile-phase B of 0.4% formic acid in a mix of 90% acetonitrile and 10% water. The flow rate was set to 230 nL/min and a 90 min gradient used (4 to 24% solvent B within 82 min, 24 to 36% solvent B within 8 min and, 36 to 100% solvent B within 1 min, 100% solvent B for 6 min before bringing back solvent B at 4% within 1 min and equilibrating for 18 min). Analysis was performed in a data-independent acquisition (DIA) mode. Full MS scans were acquired with a mass range of 375 -1250 m/z in the orbitrap, an RF lens set at 40%, and at a resolution of 120,000 (at 200 m/z). The automatic gain control (AGC) was set to a target of 4 × 10^5^, and a maximum injection time of 54 ms was applied, scanning data in profile mode. MS1 scans were followed by 41 MS2 customed windows. The MS2 scans were acquired in the Orbitrap at a resolution of 30,000 (at 200 m/z), with an AGC set to target 2 × 10^5^, for a maximum injection time of 54 ms. Fragmentation was achieved with higher energy collision induced dissociation (HCD) at a fixed normalized collision energy (NCE) of 35%. A single lock mass at m/z 445.120024 (Olsen et al., 2005) was employed. Xcalibur version 4.3.73.11 and Tune 3.4.3072.18 were used to operate the instrument. The mass spectrometry data has been deposited to the ProteomeXchange Consortium via the PRIDE partner repository (Perez-Riverol et al., 2019) with the dataset identifier PXD047504.

---

**Figure S1:**
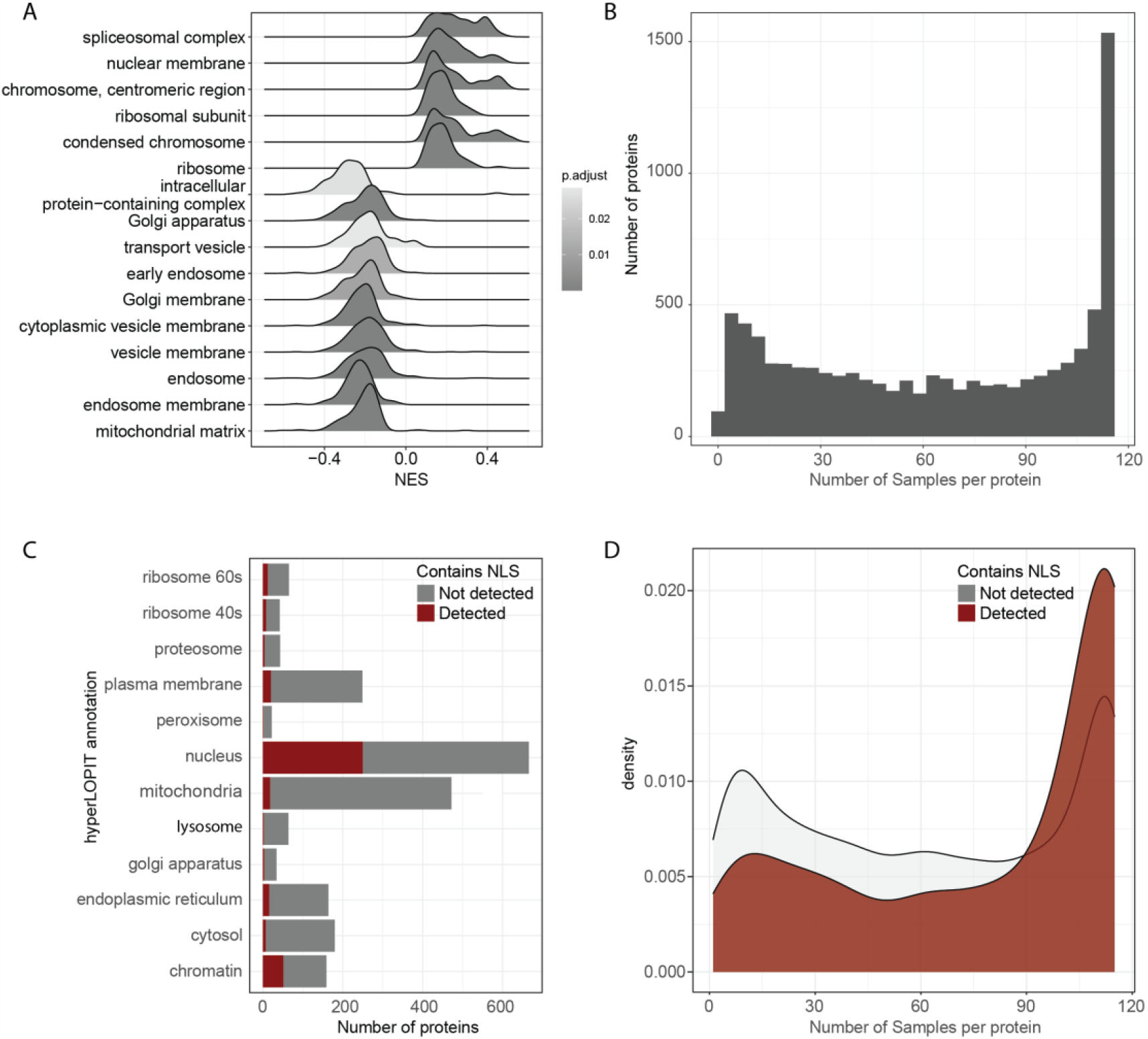
Chromatin enrichment protocol enriches for nuclear proteins and proteins with NLS a. Subcellular localisation enrichment of relative intensity of proteins found on chromatin samples. b. Number of samples each protein was detected in. c. Distribution of proteins containing an NLS, based on subcellular compartment of protein localisation in hyperLOPIT. d. Distribution of proteins containing an NLS, based on number of samples each protein was detected in.

**Figure S2:**
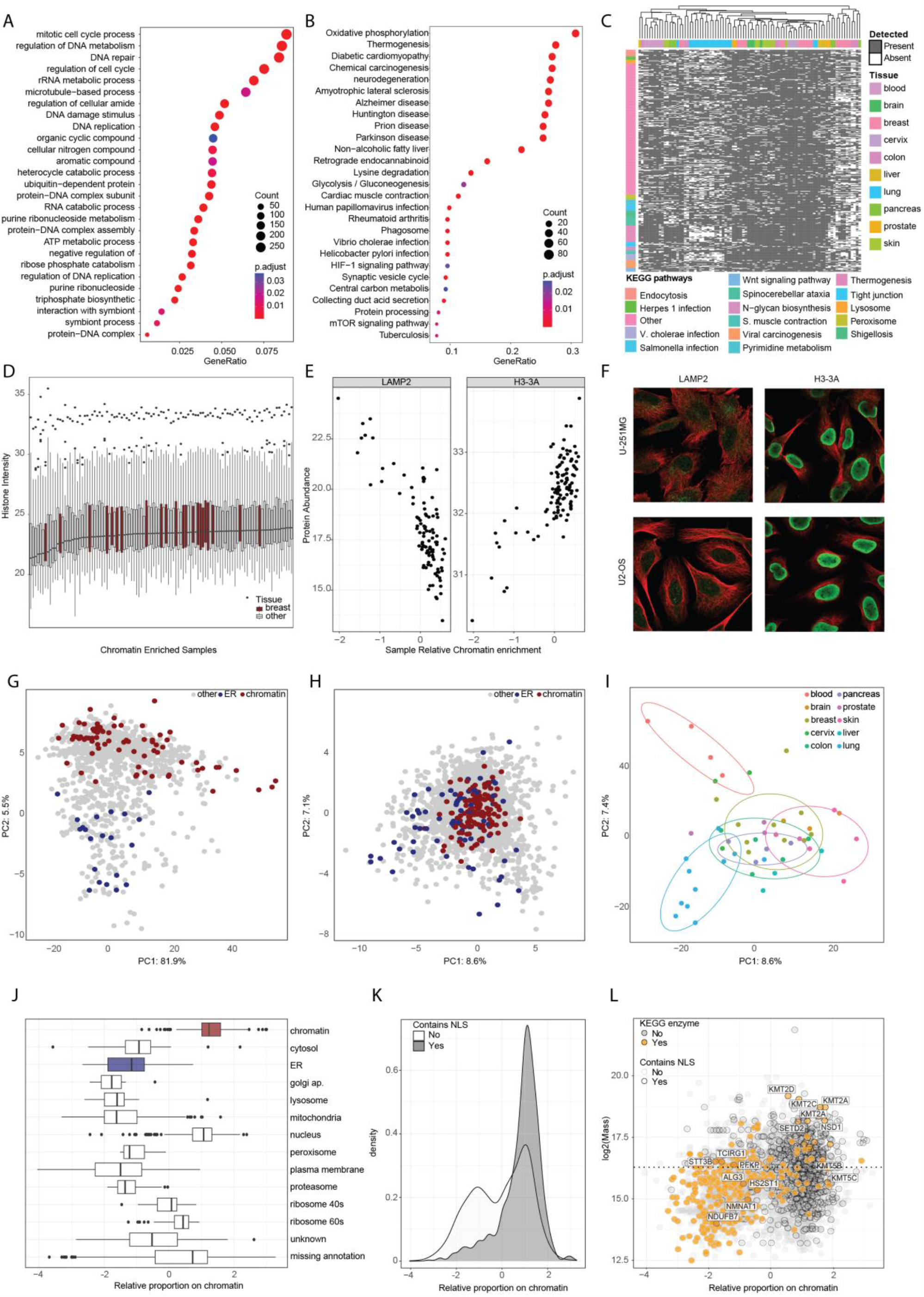
Chromatin enrichment normalization for quantitative analysis a. Biological Process enrichment of proteins found on chromatin across the majority of samples. b. KEGG pathway enrichment of proteins found on chromatin across the majority of samples. c. Clustering of chromatin proteomic samples based on detection of all KEGG proteins. d. Relative intensities of histone proteins per sample. e. Relative intensity of LAMP2 and H3-3A proteins against relative sample enrichment. f. Immunohistochemistry images in U2-OS and U-251MG cells of LAMP2 and H3-3A from HPA. g. PCA of chromatin proteins before linear regression against relative sample chromatin enrichment. h. PCA of chromatin proteins after linear regression against relative sample chromatin enrichment. i. PCA of chromatin samples after linear regression against relative sample chromatin enrichment. j. Protein slope coefficient from linear regression, based on the known subcellular protein localisation per protein according to hyperLOPIT dataset. k. Protein slope coefficient from linear regression, based on the presence of an NLS per protein. l. Protein slope coefficient from linear regression compared to protein size and presence of NLS signal.

**Figure S3:**
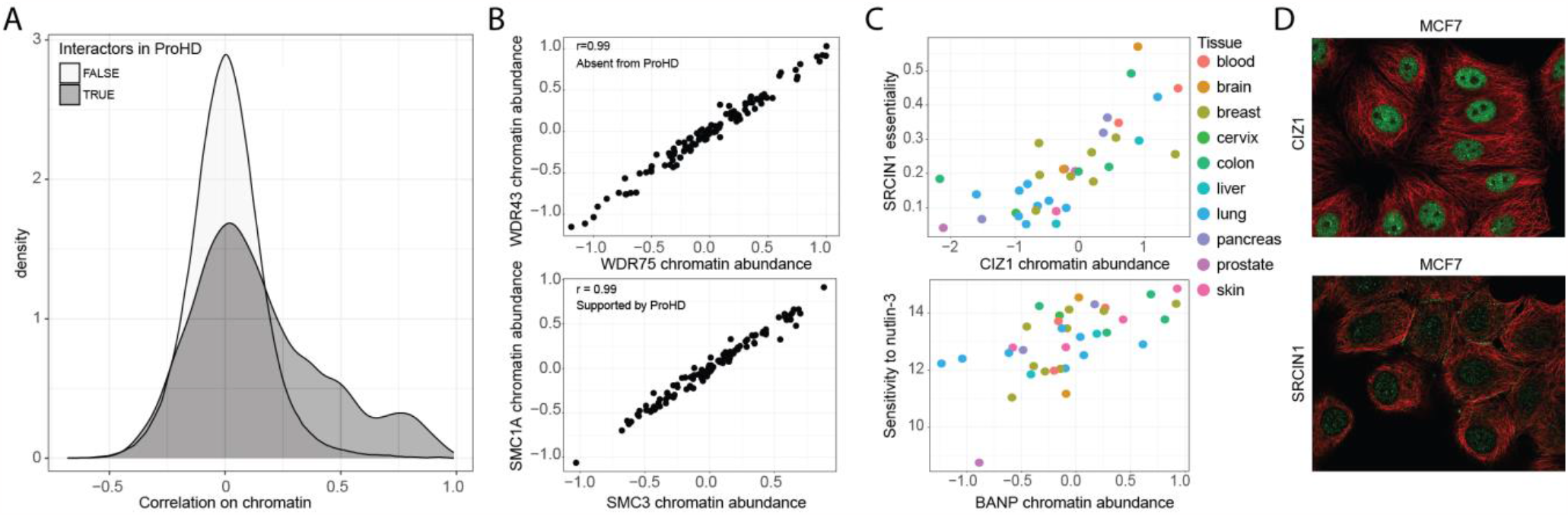
Functional relations of chromatin-localised enzymes a Protein-protein correlation, based on the known protein coregulation pairs from ProHD dataset. b. Relative protein abundance per sample for WDR75-WDR43 and SMC3-SMC1A protein pairs. c. Relative protein abundance per sample for CIZ1 against SRCIN1 gene essentiality (top) and BANP against nutlin-3 drug sensitivity. d. Immunohistochemistry images in MCF-7 cells of CIZ1 and SRCIN1 from HPA.

