## Supplementary figures and images for "Comprehensive chromatome profiling identifies metabolic enzymes on chromatin in healthy and cancer cells"

### Appendix1

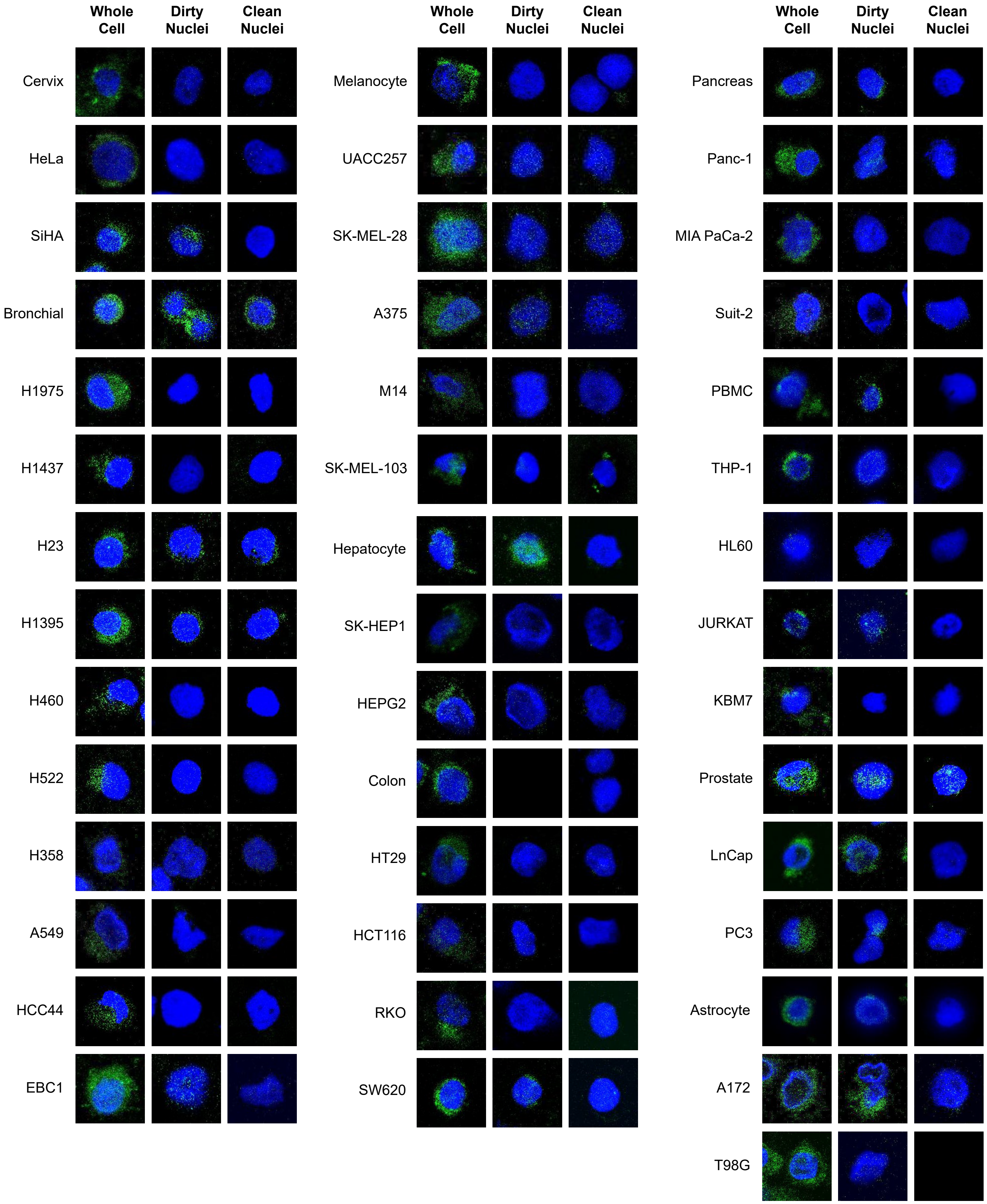

### Appendix2

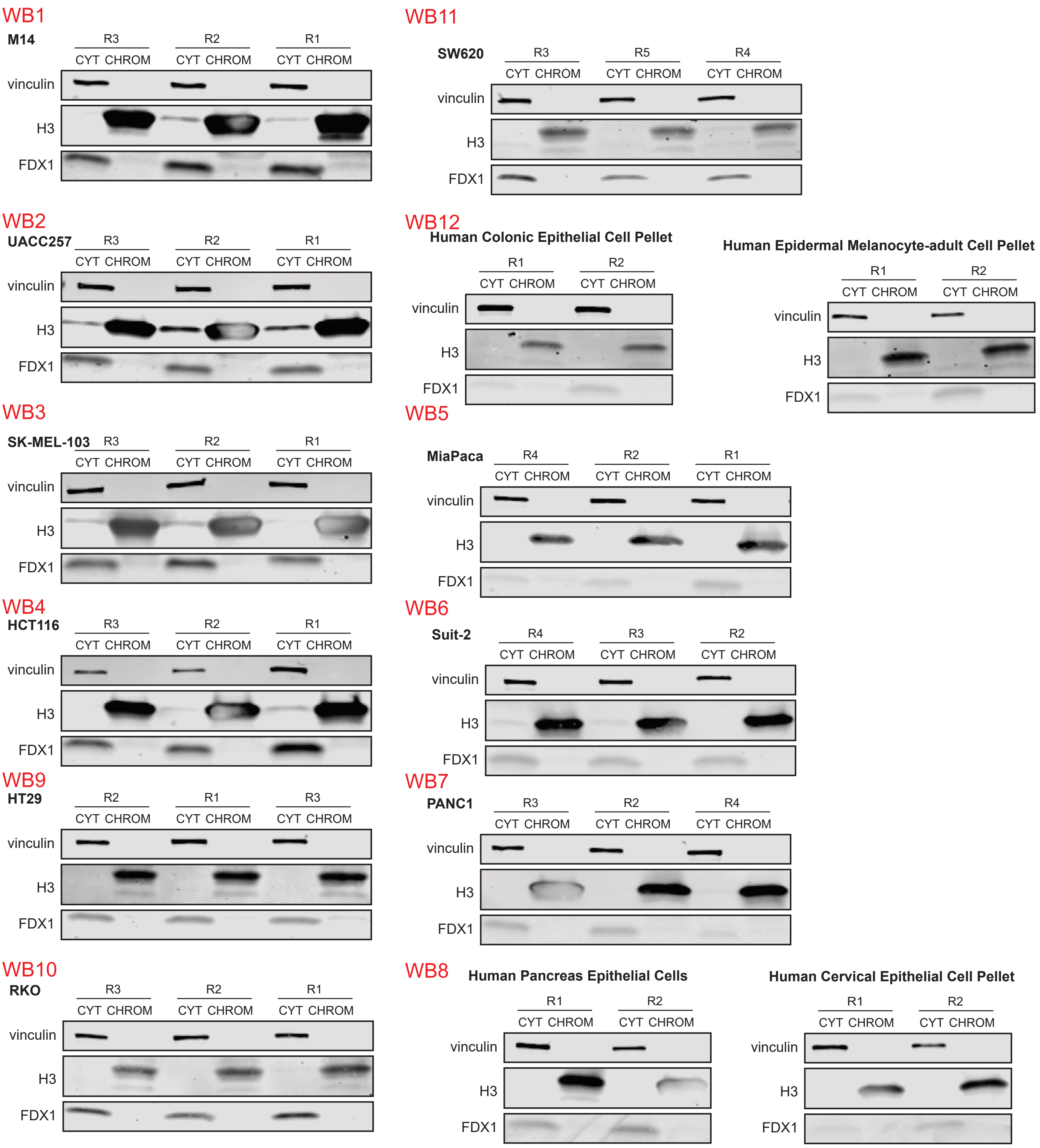

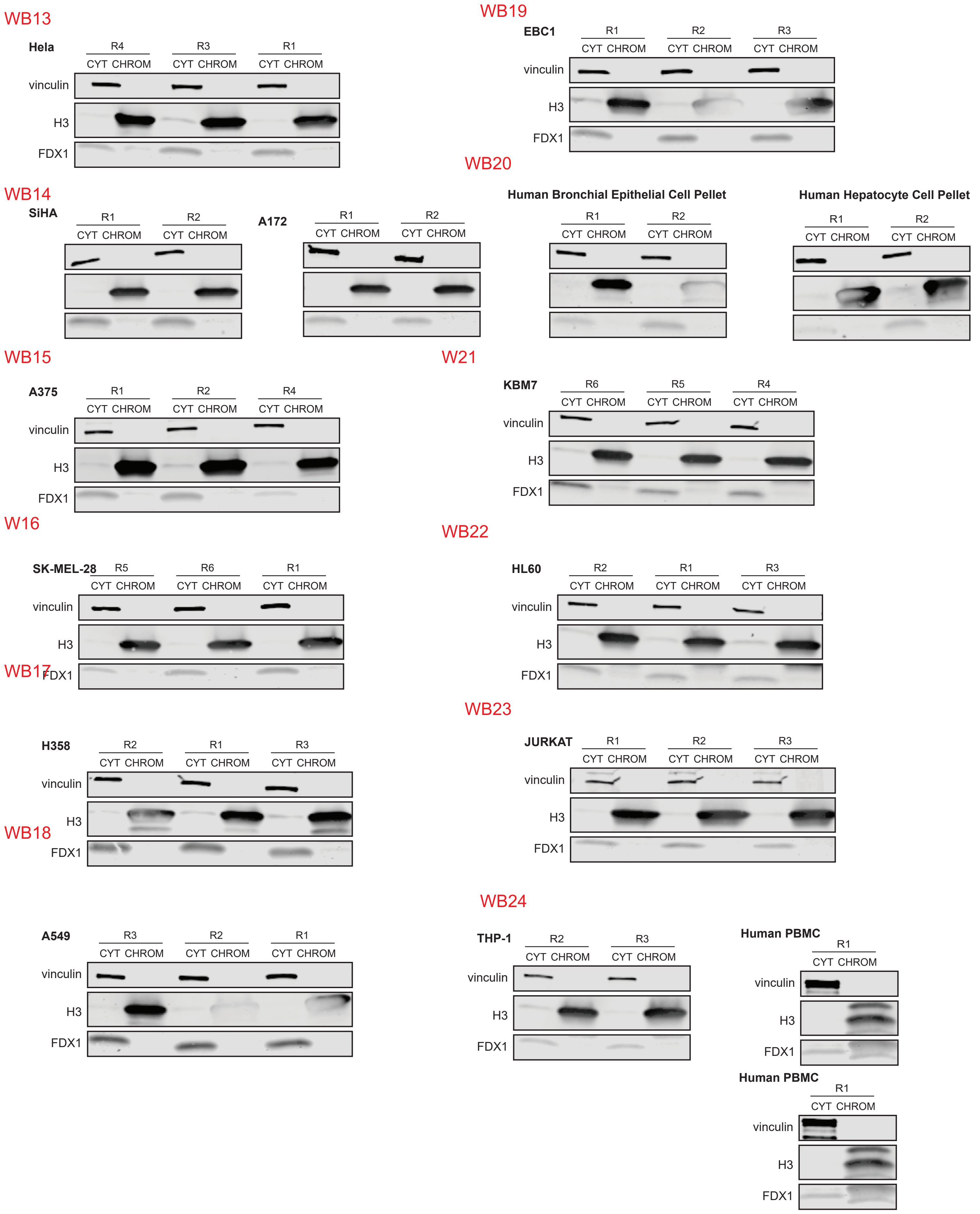

WB25

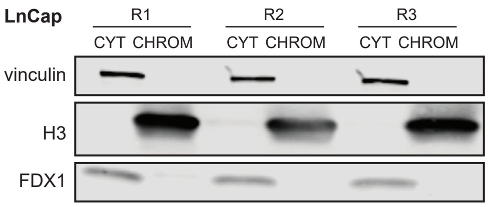


WB26

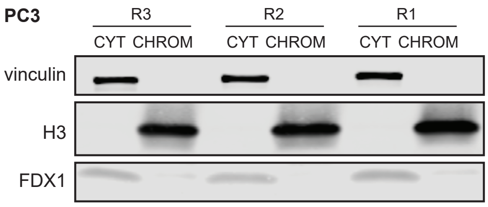


WB27

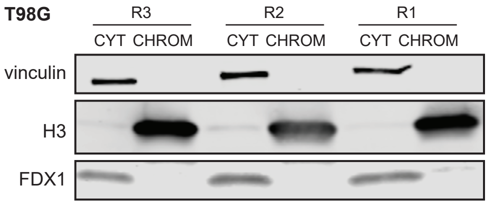


W28

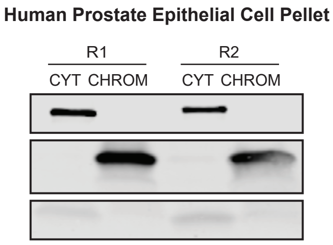


WB29

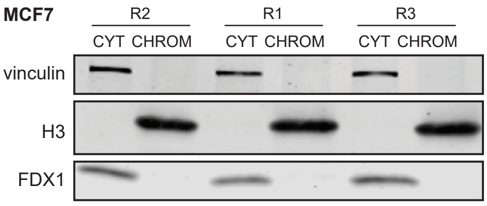


WB30

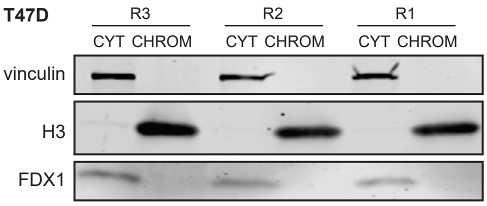


WB31

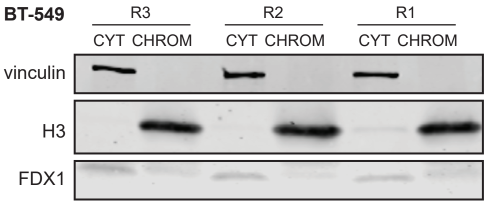


WB32

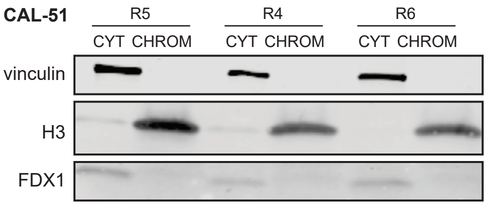


WB33

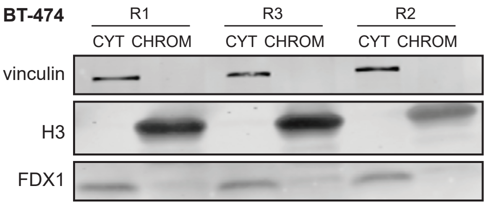


WB34

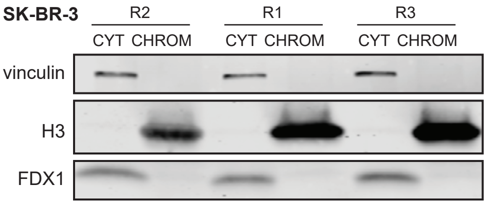


WB35

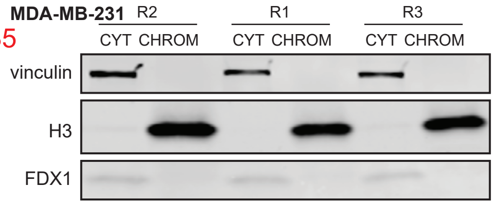


WB36

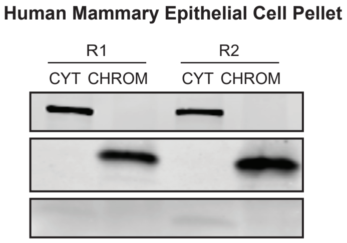

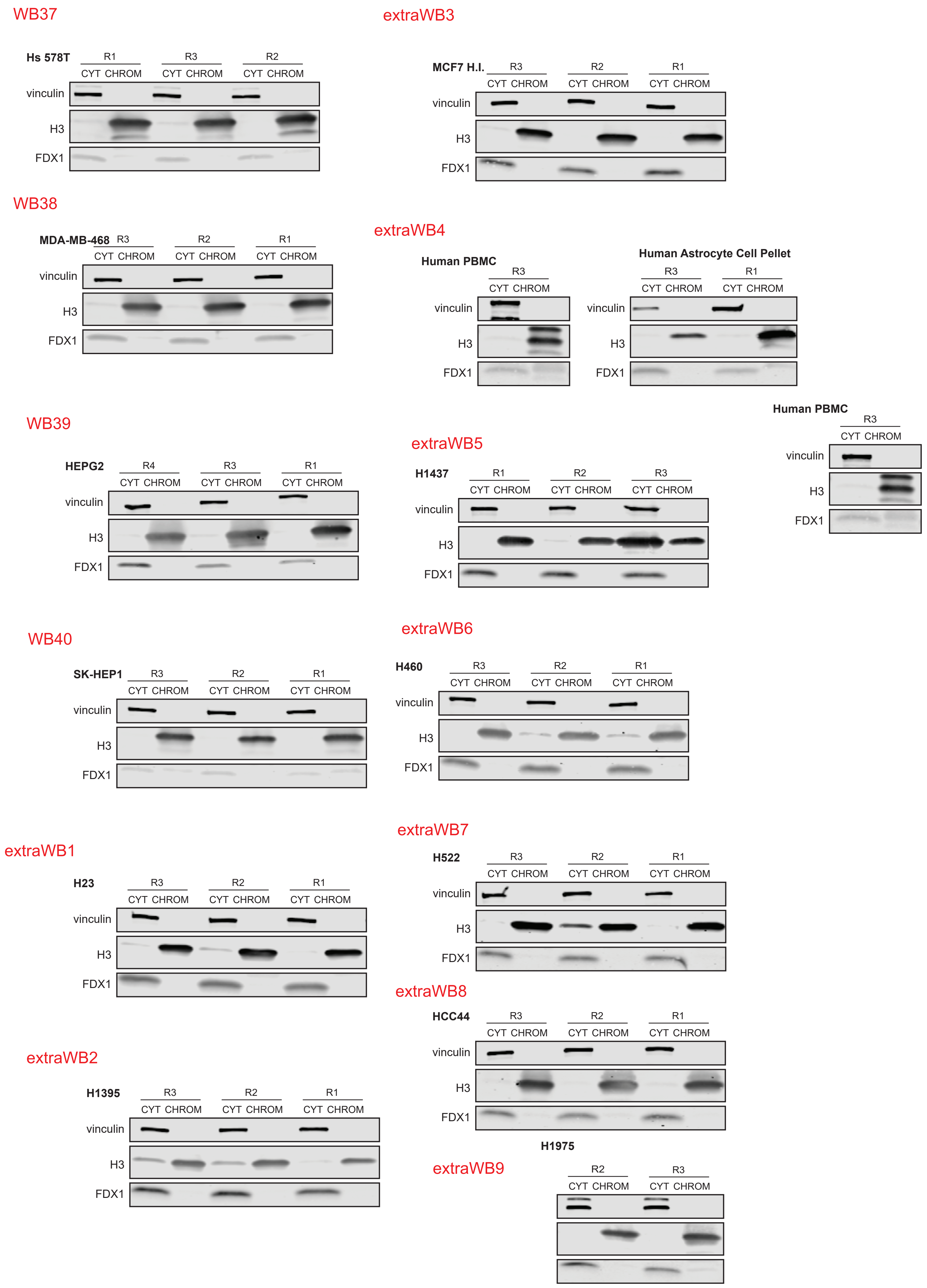
